# Phase separation drives SNARE complexes formation

**DOI:** 10.1101/2025.03.04.641570

**Authors:** Feng Chen, Qiongwei Ke, Sifan Feng, Huiyi Chen, Qin Haung, Fubin Ma, Yujie Cai, Ji Chen, Shengnan Li, Wenyan Wei, Yuanhong Sun, Xiaoping Peng, Lin Tong, Xiaocong Sun, Liehua Deng, Yan Wang, Lili Cui

## Abstract

**Background:** Neuronal exocytosis is mainly driven by the assembly of the soluble N-ethylmaleimide-sensitive factor attachment protein receptor (SNARE) complexes. However, little is known about the organization principle of the SNARE complex.

**Methods:** Protein condensates formed by SNARE complex were imaged by confocal microscope. Fluorescence recovery after photobleaching (FRAP) assay together with fusion and division assays at the cellular level and in vitro studies with purified proteins were performed to characterize the dynamic properties of protein condensates. The effect of SNARE complex phase separation on the recruitment of synaptic vesicles was detected by immunofluorescence.

**Results:** We discover that phase separation drives SNARE complex formation and aggregation; in addition, nonintrinsically disordered regions (non-IDRs) of the syntaxin1 protein is necessary for the formation of these biological condensates. Functionally, phase separation of the SNARE complex can be regulated by the major cofactors of the fusion machinery and has the ability to recruit synaptic vesicles in neurons.

**Conclusions:** Our study here establishes that phase separation is a promising way to mediate the formation and aggregation of the SNARE complex, and further identified that the non-IDRs of syntaxin1 is necessary for the phase separation of the SNARE complex. Our work answers an vital scientific question: does the SNARE complex function as multiple copies that are dispersed or clustered together to ensure sustained neurotransmitter release. In sum, phase separation provides an ideal working model for SNARE complex-mediated membrane fusion and neurotransmitter release.

## Introduction

Neuronal communication crucially relies on the exocytosis of neurotransmitters from synaptic vesicles. Neurotransmitter release is a highly regulated process that depends on the fusion of synaptic vesicles with the presynaptic membrane; this process is mediated by the soluble *N*-ethylmaleimide-sensitive factor attachment protein receptor (SNARE) complex, which is characterized by approximately 65 amino acids called SNARE motifs^1^. Neuronal SNARE complexes mainly consist of syntaxin1 and SNAP25, both of which are localized at the plasma membrane, and VAMP2 (also known as synaptobrevin 2), which is anchored to the synaptic vesicle^2^. Syntaxin1 and VAMP2 contain one SNARE motif, while SNAP25 has two SNARE motifs connected by an extended linker^3^. The fusion of the synaptic vesicular membrane with the presynaptic membrane is an energy-consuming process, and the energy needed originates from the zipper assembly of the SNARE complex^4, 5^. During membrane fusion, the three proteins form a four-helix bundle via SNARE motifs to bring the synaptic vesicle membranes and presynaptic membranes into close contact and eventually drive membrane fusion^6, 7^. Apart from these core SNARE proteins, many accessory proteins, including Sec1/Munc18 (SM) proteins ^8, 9^, Munc13^10^, synaptotagmin ^11^, and complexin ^12^, have also been shown to tightly regulate the formation and function of the SNARE complex. Although the SNARE complex is well known to govern membrane fusion, the spatial organization of the SNARE complex is unclear; in addition, whether the SNARE complex functions as a single copy or as an aggregate is unknown.

Recently, liquidLliquid phase separation (LLPS) has emerged as a ubiquitous principle that underlies the formation of membraneless compartments and has been implicated in many biological processes, ranging from gene expression to signal transduction^13, 14^. LLPS has also been described as an important method for the assembly of protein networks. Recently, an increasing number of observations have indicated that synaptic transmissions in both presynaptic boutons and postsynaptic densities are likely such biological condensates. Studies have provided substantial evidence that phase separation underlies the organization of the presynaptic membrane active zone^15, 16^, the clustering and maintenance of synaptic vesicles^17, 18^, the exocytosis and endocytosis of synaptic vesicles^19, 20^, the formation of postsynaptic density assemblies^21^, and even the short-distance vesicle transport^22^. However, whether the assembly and clustering of the SNARE complex (the core mechanism of synaptic transmission) are mediated by phase separation is currently unknown.

In the present study we discover that phase separation drives SNARE complex formation and aggregation through fluorescence recovery after photobleaching (FRAP) assay and droplet fusion or division assays at the cellular level and in vitro studies with purified proteins. In addition, we identify that syntaxin1 syntaxin1 (210-288 amino acids) is necessary for the phase separation of SNARE complex. Functionally, we found that SNARE complex phase separation has the ability to recruit synaptic vesicles in rat neurons. Together, our findings reveal that phase separation is a promising way to mediate the formation and aggregation of the SNARE complex, and possility an important foundation for future directions on the mechanisms underlying membrane fusion, neurotransmission, and even neurological disorders associated with the SNARE complex.

## Materials and methods

### Cell culture, transfection, and imaging

Human embryonic kidney (HEK)-293T cells were cultured in Dulbecco’s modified Eagle’s medium (DMEM) (Gibco, 8122669) supplemented with 10% fetal bovine serum (FBS) (Invigentech, A6903FBS-500) and 1% penicillinLstreptomycin (Gibco, 15140-122) at 37°C in a 5% CO_2_ incubator. For primary neuronal cultures, we dissected and cultured mouse hippocampal or cortical neurons from C57BL/6 mice on the P1, as previously described^23^. The dissociated neurons were seeded on 35-mm poly-D-lysine (PDL)-coated glass-bottom dishes and maintained in Neurobasal-A medium (Gibco, 21103-049) supplemented with B-27 (Gibco, 17504-044). After 5 days in vitro, the cells were transfected with the corresponding combinations of these plasmids (pCMVPuro-VAMP2-mCherry, pCMVPuro-syntaxin1-ECFP, pCMVPuro-SNAP25-EGFP, pCMVPuro-VAMP2-EGFP, pCMVPuro-STX1A-EGFP, pCMVPuro-Munc18-1, and pCMVPuro-Munc13-1) at ∼ 60% confluence by using Lipofectamine 3000 (Invitrogen, L3000015) according to the manufacturer’s instructions. After 24 h, live-cell imaging was performed directly (Olympus FV3000 Laser Confocal Microscope), or imaging was performed after fixation with 4% paraformaldehyde for 15 minutes. Rats study was conducted in accordance with the protocol approved by the Laboratory Animal Ethics Committee of Guangdong Medical University.

### Fluorescence recovery after photobleaching (FRAP) assay

FRAP assays in HEK-293T cells were performed on an Olympus FV3000 Laser Confocal Microscope with a 60x oil lens immersion objective. FRAP was conducted using 488 nm laser beams at 2%-20% power for bleaching. The distribution of the fluorescence intensity within the photomanipulated droupts was recorded over time. The intensity at the maximalprebleach point was normalized to 100%. The time immediately after the bleaching event was set to 0 seconds.

### In vitro phase separation experiments

The recombinant proteins VAMP2-mcherry (GenScrip, C798T461G0-8), SNAP25-EGFP (GenScrip, C798T461G0-2), and STX1A-ECFP (GenScrip, C798T461G0-5) at the indicated concentrations were mixed in test tubes. At a volume ratio of 1:1:1, different concentrations of PEG-8000 (Diamond, A100159-0500) and different concentrations of NaCl were added to the tubes. The solution was mixed well and incubated at room temperature for 10 minutes. Protein droplets were imaged on glass-bottom dishes by using an Olympus FV3000 laser confocal microscope.

### Droplet fusion or division assay

For phase-separated droplets formed in living HEK-293T cells, an Olympus FV3000 laser confocal microscope with a 60x oil lens was used to take long-term images. After the relevant cells were identified, images were taken first, and then all lasers were turned off to avoid fluorescence quenching. Five minutes later, the laser was turned on, and fluorescence images were obtained. The above steps were repeated to record the fusion and division of phase separation droplets. For live imaging of recombinant protein fusion or division in condensed droplets, the total volume of each mixture was 150 μL. Images of the droplets were taken every 10 seconds to observe the fusion and division of the droplets.

### Immunofluorescence and colocalization

The cells were washed with ice-cold PBS and fixed with 4% paraformaldehyde for 30 min at room temperature. The cells were permeabilized with 0.1% Triton X-100 (Beyotime, ST797) for 5 min, blocked with 5% horse serum for 1 h, and then incubated with primary antibodies (Proteintech, rabbit anti-CD63, 25682-1-AP, 1:200; Proteintech, rabbit anti-TSG101, 28283-1-AP, 1:200; Proteintech, rabbit anti-synaptophysin, 17785-1-AP, 1:300) overnight at 4°C. The cells were washed three times with PBS and incubated with fluorescently labeled secondary antibodies (Abcam, goat anti-rabbit IgG H&L (Alexa Fluor® 594), AB150080, 1:10000) for 1 h at room temperature. The colocalization of protein droplets with membrane organelles, exosomal proteins, and synaptic vesicles was detected by an Olympus FV3000 laser confocal microscope with a 60× oil objective lense.

### Protein expression and purification

All purified proteins in this study were expressed in a pET30a vector. Constructs encoding syntaxin1, VAMP2 and SNAP25 were fused with the ECFP, mCherry, or EGFP coding sequence and subcloned and inserted into a pET30a expression vector with a His tag at the N-terminus. The fusion proteins were overexpressed in BL21 (DE3) *E. coli* cells. Protein expression was induced by the addition of 50 mM isopropyl β-D-1-thiogalactopyranoside (IPTG), and the cells were further cultured at 37°C for 16 h and then lysed with an ultrasonicator on ice. The samples were then centrifuged at 30,000 × g for 1 h at 4°C, after which the cell debris was removed. The supernatant was incubated with glutathione agarose beads at 4°C for 2 h to pull down the proteins. Proteins were further eluted with reduced glutathione in lysis buffer. The eluted protein was purified via high-performance liquid chromatography. The percentage of purified proteins was greater than 75%, as determined by SDSLPAGE under reducing conditions, and the protein concentration was calculated by using BSA as a standard. The final purified proteins were then equilibrated with storage buffer (150 mM NaCl, 10% glycerol, 50 mM Tris-HCl, pH 8.0) and stored at −80°C until further use.

### Statistical analysis

Statistical tests were performed using GraphPad Prism 9.0. All the data are presented as the mean ± standard error of the mean (SEM). All the experiments were repeated independently at least 3 times. Comparisons between two groups were performed by Student’s t tests. Statistical significance was set at *p* < 0.05 (\**p* < 0.05, \*\**p* < 0.01, and \*\*\**p* < 0.001). All of the statistical details can be found in the figure legends.

## Results

### The SNARE complex forms protein condensates in vitro

Intrinsically disordered regions (IDRs) are well recognized to drive phase separation. To determine whether the SNARE proteins sequence in the monomeric form contains IDRs, we performed bioinformatics analysis using PONDR algorithms (http://pondr.com/) to predict the presence of IDRs in the SNARE proteins and found that these proteins harbor extensive IDRs. The specific sequences are VAMP2 (residues 1-81) (Fig. 1a), SNAP25 (residues 1-81 and 137-206) (Fig. 1b), and syntaxin1 (residues 1-209) (Fig. 1c). To assess whether SNARE proteins can phase separate into liquid-like condensates, enhanced green fluorescent protein (EGFP)-tagged VAMP2 (VAMP2-EGFP), SNAP25 (SNAP25-EGFP) and syntaxin1 (syntaxin1-EGFP) plasmids were cotransfected into HEK-293T cells for 24 h. Strong protein aggregation was observed in cells cotransfected with equal amounts of VAMP2 and syntaxin1 plasmids, SNAP25 and syntaxin1 plasmids, or VAMP2 and SNAP25 and syntaxin1 plasmids (Fig. S1a and b). However, when the cells were cotransfected with VAMP2 and SNAP25, no droplets were observed. Instead, the green fluorescence was uniformly distributed in the cytoplasm. When the cells were transfected with EGFP as a control, no droplets were detected. To further evaluate whether the droplets were formed by these SNARE proteins, mCherry-tagged VAMP2 (VAMP2-mCherry), SNAP25-EGFP and ECFP-tagged syntaxin1 (syntaxin1-ECFP) constructs labeled with different colors of fluorescence were cotransfected into HEK-293T cells. Consistent with the above results, no droplets were detected in the cells cotransfected with VAMP2-mCherry and SNAP25-EGFP (Fig. 1d), while droplets were detected in the cells cotransfected with mCherry-VAMP2 and syntaxin1-ECFP (Fig. 1e), SNAP25-EGFP and syntaxin1-ECFP (Fig. 1f), or VAMP2-mCherry and SNAP25-EGFP and syntaxin1-ECFP (Fig. 1g). Fluorescence colocalization analysis revealed that the different fluorescence almost completely overlapped (Fig. 1h-j), suggesting that the droplets were formed by these SNARE proteins. These results indicate that SNARE proteins can potentially phase separate into condensates and that syntaxin1 is necessary for condensate formation. In addition, SNARE complex droplets did not colocalize with membrane-bound organelles, such as lysosomes and mitochondria (Fig. S2a), while the droplets colocalized with the endoplasmic reticulum to some extent (Fig. S2b). Furthermore, the immunofluorescence results showed that the droplets were colocalized with neither vesicle-associated proteins (Rab5) (Fig. S2c) nor exosomal proteins (CD63 and TSG101) (Fig. S2d), which further confirmed that these droplets were not exosomes or vesicles. Collectively, these results indicate that the SNARE complex forms protein condensates in cells.

**Fig. 1.**
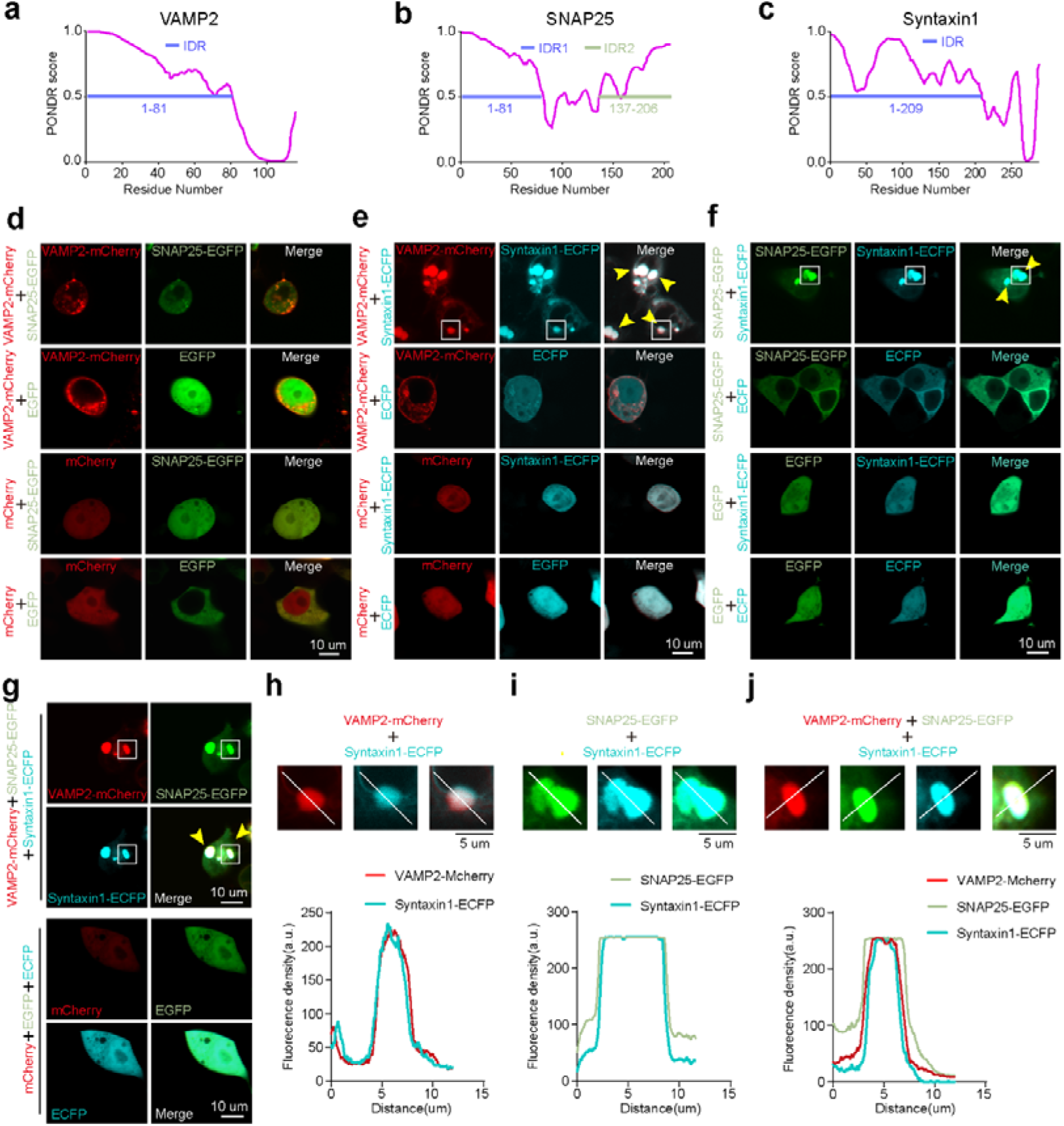
The SNARE complex forms protein condensates in vitro. Relative disorder scores of the full-length VAMP2 (**a**), SNAP25 (**b**) and syntaxin1 (**c**) proteins as predicted by the PONDR algorithm (http://pondr.com/). The amino acid positions are presented on the X-axis, and the PONDR scores are shown on the Y-axis. **d** Representative images of HEK-293T cells transfected with 1 µg/ml VAMP2-mCherry and SNAP25-EGFP, VAMP2-mCherry and EGFP, mCherry and SNAP25-EGFP or mCherry and EGFP plasmids for 24 h. (scale bar, 10 µm). **e** Representative images of HEK-293T cells transfected with 1 µg/ml VAMP2-mCherry and syntaxin1-ECFP, VAMP2-mCherry and ECFP, mCherry and syntaxin1-ECFP or mCherry and ECFP plasmids for 24 h. (scale bar, 10 µm). **f** Representative images of HEK-293T cells transfected with 1 µg/ml of SNAP25-EGFP and syntaxin1-ECFP, SNAP25-EGFP and ECFP, EGFP and syntaxin1-ECFP or EGFP and ECFP plasmids for 24 h. (scale bar, 10 µm). **g** Representative images of HEK-293T cells transfected with 1 µg/ml of VAMP2-mCherry and SNAP25-EGFP and syntaxin1-ECFP or mCherry and EGFP and ECFP plasmids for 24 h. (scale bar, 10 µm). **h**-**j** In the magnified images, the lines indicate the distribution of the fluorescence intensity of the droplets in the white squares in Fig. 1. e, f and g, respectively. Scale bar, 5 μm.

### The SNARE complex undergoes phase separation in vitro

Internal diffusion, diffusion across boundaries, fusion and fission are important criteria for phase separation condensates. Thus, FRAP experiments were performed to analyze the dynamics of the proteins inside the condensates and characterize the condensates in more detail. The results showed that the components of VAMP2-EGFP and syntaxin1-EGFP (Fig. 2a), SNAP25-EGFP and syntaxin1-EGFP (Fig. 2b), and VAMP2-EGFP, SNAP25-EGFP and syntaxin1-EGFP (Fig. 2c) recovered up to approximately 40% of their initial concentration within 1 min, indicating that the protein constituent droplets were in a liquid-like state and could transfer between the condensate droplets and the surrounding dilute solution. To eliminate interference from GFP, VAMP2-mCherry and syntaxin1-ECFP (Fig. S3a), SNAP25-EGFP and syntaxin1-ECFP (Fig. S3b), and VAMP2-mCherry, SNAP25-EGFP and syntaxin1-ECFP (Fig. S3c) were overexpressed in HEK-293T cells, which were subjected to FRAP experiments. In addition, a slow light recovery process was observed, and the droplets tended to slowly fuse with the surrounding droplets; the droplets also split into two smaller droplets, revealing the dynamic nature of the droplets in living mammalian cells (Fig. 2d-f). 1,6-Hexanediol (1,6-HD) is a useful tool for inhibiting phase separation putatively by disrupting weak hydrophobic proteinLprotein interactions^24^. To further characterize these droplets, we evaluated the effect of 1,6-HD on the condensates of the SNARE complex, and the puncta were exposed to different concentrations (0%, 2%, and 5%) of 1,6-HD for varying durations (0, 2, 5, 10, and 20 min). We found that the puncta of the SNARE complex drastically dissolved with increasing 1,6-HD exposure time and with increasing 1,6-HD concentration (Fig. S4a, b), indicating that 1,6-HD exerted time- and dose-dependent effects on the biomolecular condensates of the SNARE complex. These criteria demonstrate the phase separation properties of SNARE complex droplets in cells.

**Fig. 2.**
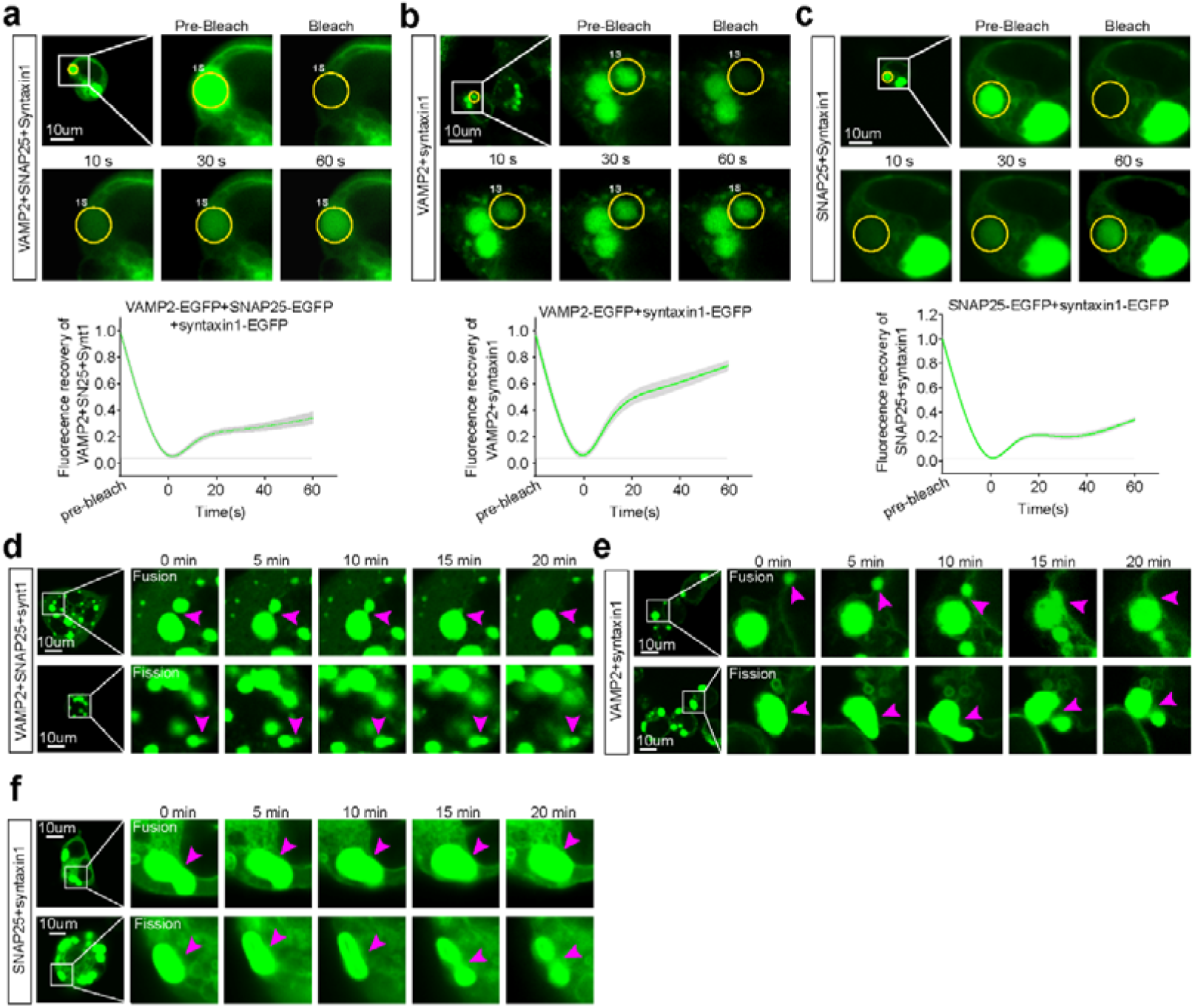
The SNARE complex undergoes phase separation in vitro. Representative images and quantification of fluorescence intensity from live-cell imaging of VAMP2-EGFP, SNAP25-EGFP and syntaxin1-EGFP constructs (**a**), VAMP2-EGFP and syntaxin1-EGFP constructs (**b**), and SNAP25-EGFP and syntaxin1-EGFP constructs (**c**) overexpressed in HEK-293T cells (1 µg/ml for 24 h) before and after photobleaching. The yellow dashed lines indicate photobleached regions. Scale bar: 10 μm. Six droplets were used for FRAP analysis. Representative live-cell images showing dynamic fusion and fission events of VAMP2-EGFP and SNAP25-EGFP and syntaxin1-EGFP puncta (**d**), VAMP2-EGFP and syntaxin1-EGFP puncta (**e**), and SNAP25-EGFP and syntaxin1-EGFP puncta (**f**) in HEK-293T cells transfected with each of these constructs (1 µg/ml for 24 h). Scale bar: 10 μm. The data are presented as the mean ± SEM; \**p* < 0.05; \*\**p* < 0.01; \*\*\**p* <0.001.

### Phase separation of the SNARE complex in vitro

To confirm that the results were not compromised by the complicated cellular environment of cells, we purified recombinant VAMP2-mCherry, SNAP25-EGFP and syntaxin1-ECFP fusion proteins. When we combined these proteins (VAMP2-mCherry, syntaxin1-ECFP and SNAP25-EGFP), they spontaneously aggregated to form dose-dependent visible condensates within 10 minutes in the presence of a macromolecular crowder, polyethylene glycol (PEG-8000, 10% w/v), and salt (1 M NaCl). Syntaxin1-ECFP/SNAP25-EGFP complex puncta first appeared at a concentration of approximately 2 μM, and the puncta number significantly increased with increasing amounts of protein (Fig. 3a and b). The same trend was observed when the VAMP2-mCherry and syntaxin1-ECFP recombinant proteins were mixed (Fig. 3c and d) or when all three SNARE recombinant proteins were mixed (Fig. 3e and f). These data suggest that the formation of SNARE complex phase separation is dose dependent during in vitro studies with purified proteins. It is well known that the ionic environment and crowding agent significantly impact phase separation. To determine the effect of salt on SNARE complex phase separation, NaCl at different concentrations (0.1, 0.5, 1.0, 2.0, and 5.0 M) was added to the protein complex, and the results showed that a low salt environment (< 1.0 M) promoted phase separation, which may be explained by a decrease in intramolecular electrostatic interactions among the SNARE proteins. A high concentration of salt (> 2.0 M) drastically inhibited the formation of phase separation in the SNARE complex, as the droplets disappeared when the concentration of NaCl reached 5 mol; these results suggest that the SNARE complex condenses in a salt-sensitive manner (Fig. S5a-f). Unlike salt, high concentrations of the crowding agent may promote the phase separation of the SNARE complex, as the number of droplets rapidly increased as the concentration of PEG-8000 increased (Fig. S6a-f). Corresponding with the results of the intracellular experiments, we also observed that the two droplets coalesced to form a larger structure, and one droplet can also split into two smaller condensates (Fig. 3g-i). Moreover, the formation of SNARE protein droplets almost disappeared upon exposure to 1,6-HD, indicating sensitivity to phase separation inhibitors in vitro (Fig. S7**a-c**). Taken together, these data indicate that the SNARE complex undergoes phase separation in vitro.

**Fig. 3.**
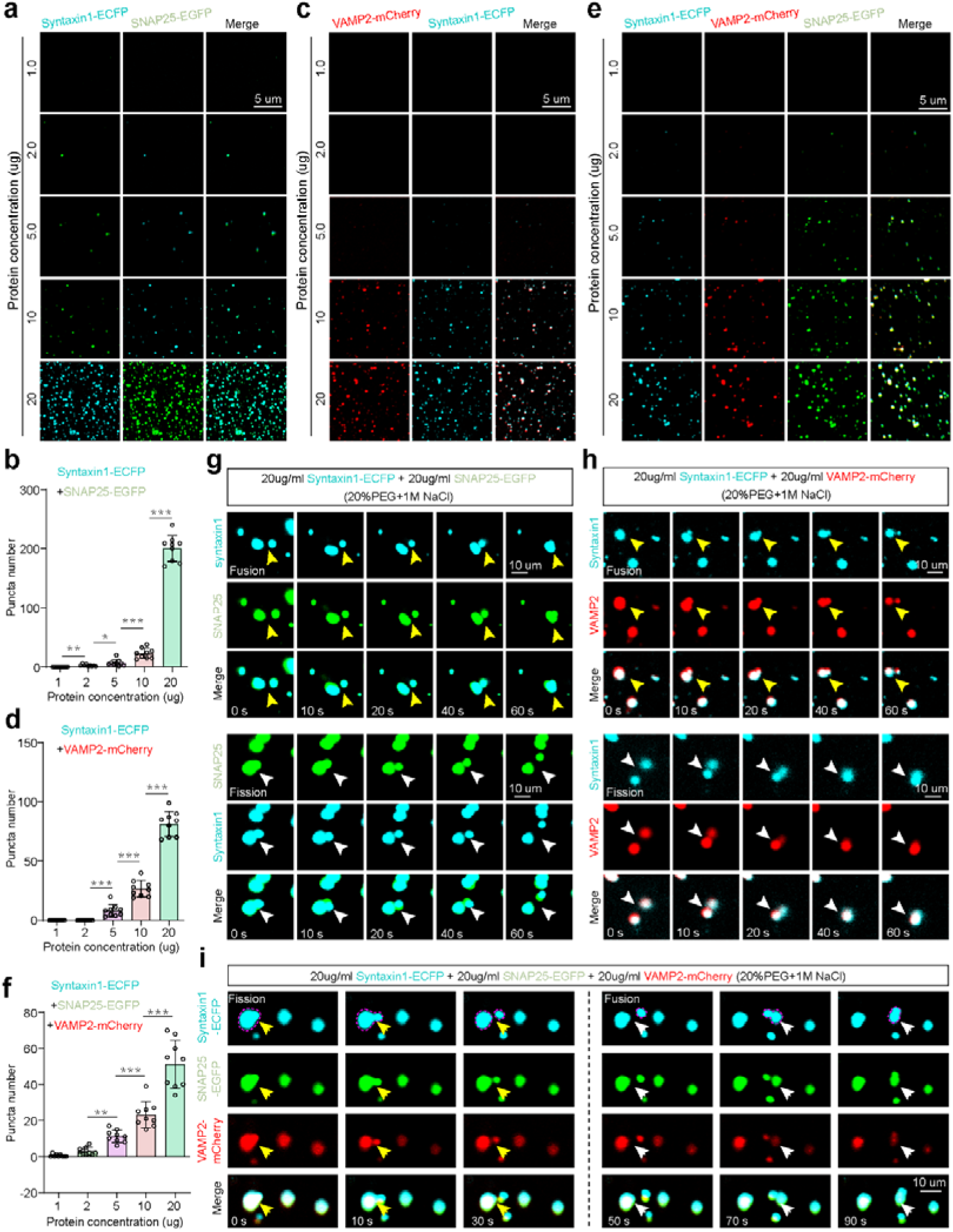
The SNARE complex undergoes phase separation in a cell-free system. **a**, **b** Representative images and quantification of syntaxin1-ECFP and SNAP25-EGFP fusion proteins at different concentrations (1.0, 2.0, 5.0, 10.0, 20.0 µg) in buffer containing crowding reagent (PEG-8000, 10% w/v) and salt ions (1 M NaCl). Scale bar, 5 μm. **c**, **d** Representative images and quantification of VAMP2-mCherry and syntaxin1-ECFP and SNAP25-EGFP fusion proteins at different concentrations (1.0, 2.0, 5.0, 10.0, 20.0 µg) in buffer containing crowding reagent (PEG-8000, 10% w/v) and salt ions (1 M NaCl). Scale bar, 5 μm. **e**, **f** Representative images and quantification of syntaxin1-ECFP, VAMP2-mCherry and SNAP25-EGFP fusion proteins at different concentrations (1.0, 2.0, 5.0, 10.0, 20.0 µg) in buffer containing crowding reagent (PEG-8000, 10% w/v) and salt ions (1 M NaCl). Scale bar, 5 μm. Fusion (yellow arrowheads) and fission (white arrowheads) events of droplets formed by the syntaxin1-ECFP and SNAP25-EGFP fusion proteins (**g**), and syntaxin1-ECFP and VAMP2-mCherry fusion proteins (**h**), and syntaxin1-ECFP, VAMP2-mCherry and SNAP25-EGFP fusion proteins (**i**). Scale bar: 10 μm. Nine fields of view were randomly selected for statistical analysis. The data are presented as the mean ± SEM; \**p* < 0.05; \*\**p* < 0.01; \*\*\**p* <0.001.

### Syntaxin1 is necessary for the phase separation of the SNARE complex

As demonstrated above, the syntaxin1 protein is necessary for the SNARE complex to undergo phase separation. To further clarify this idea, we overexpressed the syntaxin1 plasmid in HEK-293T cells cotransfected with the mCherry-VAMP2 and SNAP25-EGFP plasmids to observe its effect on SNARE complex phase separation, and the results showed that the dose (0, 0.1, 0.5, 1.0, 2.0, or 3.0 µg/ml) significantly increased the droplets size of the SNARE complex (Fig. 4**a**, Fig. S8**a**). Consistent with the results obtained at the cellular level, we observed protein dose (0, 1.0, 5.0, 10.0, 2.0, and 30.0 µg/ml)-dependent SNARE complex phase separation based on in vitro studies with purified proteins (Fig. 4**b**, Fig. S8**b**). To further confirm this phenomenon, we transfected mCherry-VAMP2 and SNAP25-EGFP constructs into HEK-293T cells, and no protein condensates were observed after 24 hours (Fig. 4**c**, **d**). However, obvious protein droplets were found when the syntaxin1 construct was further overexpressed in cells for another round of 24 hours (Fig. 4**e**, **f**). Taken together, these findings reveal that syntaxin1 is an essential factor for SNARE complexes to undergo phase separation.

**Fig. 4.**
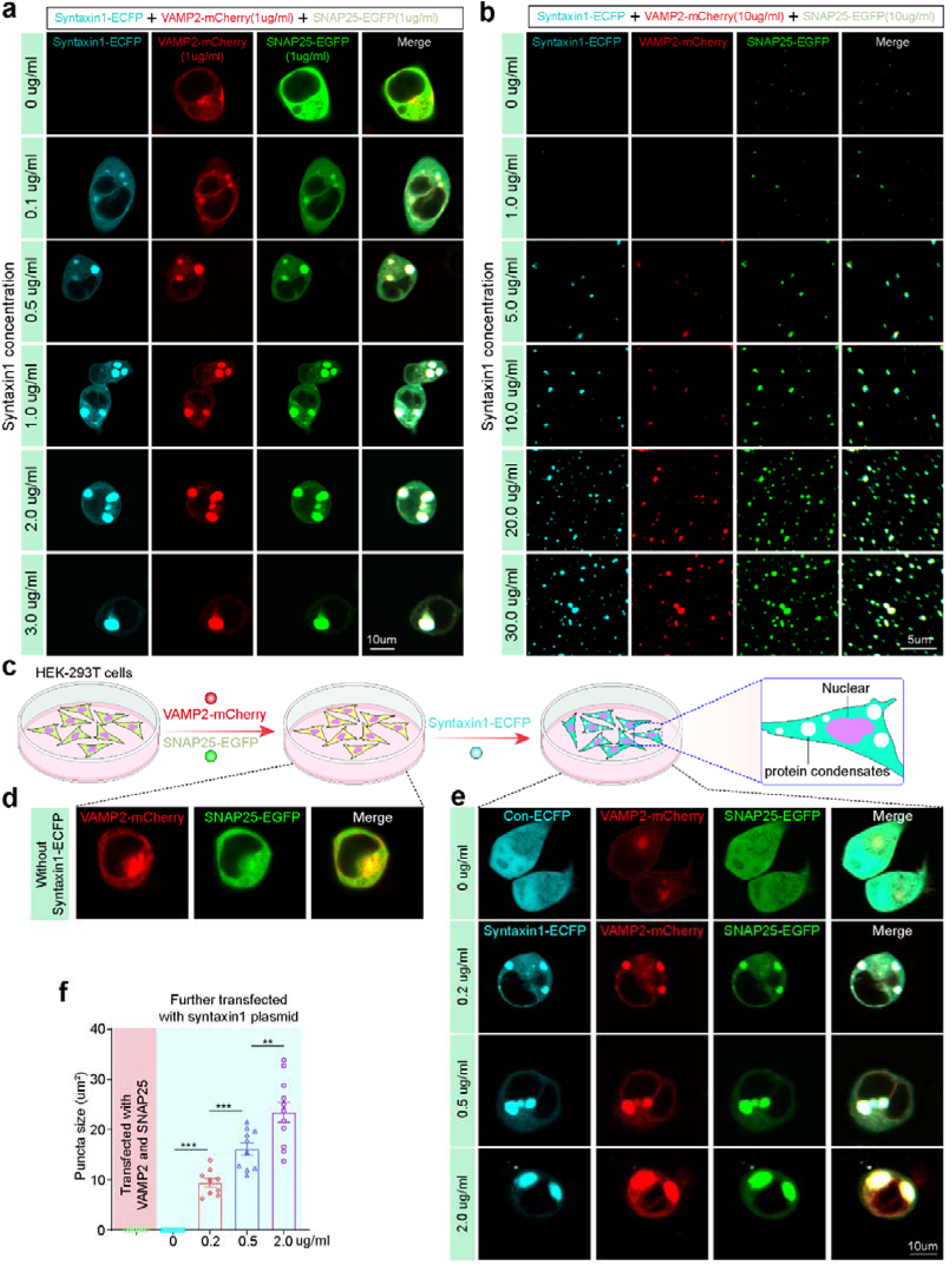
Syntaxin1 is necessary for the phase separation of the SNARE complex. **a** HEK-293T cells were overexpressed with VAMP2-mCherry (1 µg/ml), SNAP25-EGFP (1 µg/ml) and increasing amounts of syntaxin1-ECFP constructs (0, 0.1, 0.5, 1.0, 2.0, or 3.0 µg/ml) for 24 h, and representative images were obtained by confocal microscopy. Scale bar: 10 μm. **b** VAMP2-mCherry (1 µg/ml), SNAP25-EGFP (1 µg/ml) and increasing amounts of syntaxin1-ECFP fusion proteins were subjected to buffer containing crowding reagent (PEG-8000, 10% w/v) and NaCl (1.0 mol) for 10 minutes, and representative images were obtained by confocal microscopy. **c** Flow chart showing the experimental approach used to clarify the effect of syntaxin1 on the phase separation of the SNARE complex. **d, e** HEK-293T cells were cotransfected with VAMP2-mCherry (1 µg/ml) and SNAP25-EGFP (1 µg/ml) constructs for 24 h, and then the cells were further transfected with increasing amounts of syntaxin1-ECFP constructs (0, 0.2, and 2.0 µg/ml) for 24 h. Representative images were obtained by confocal microscopy. Scale bar: 10 μm. **f** Quantification of **d** and **e**; at least six cells were randomly selected. The data are presented as the mean ± SEM. \**p* < 0.05; \*\**p* < 0.01; \*\*\**p* <0.001.

### The non-IDRs of syntaxin1 is necessary for the phase separation of the SNARE complex

To further identify the key domains in SNARE proteins that are needed for phase separation, a series of truncated different fluorescent protein-labeled mCherry-VAMP2 (residues 1-81 and residues 82-118), SNAP25-EGFP (residues 1-81 and 82-136 and residues 137-206), and syntaxin1-ECFP (residues 1-209 and residues 210-288) plasmids were constructed according to the IDR region of the proteins. The formation of SNARE complex condensates was not affected by deletion of the IDRs or non-IDRs of the VAMP2 protein (Fig. 5**a**) and SNAP25 protein (Fig. 5**b**). Unexpectedly, the removal of the IDR of the syntaxin1 protein did not affect SNARE complex phase separation; in contrast, the deletion of the non-IDR did not cause droplets to form (Fig. 5**c**). On the other hand, no condensates were detected when the IDRs of syntaxin1 together with VAMP2 and the truncated SNAP25 plasmids were used in HEK-293T cells (Fig. 5**d**). Taken together, these data indicate that the non-IDRs of the syntaxin1 protein are the key region responsible for the phase separation of the SNARE complex. Next, we tried to identify the key amino acids in the non-ID region of syntaxin1 that are responsible for the phase separation of the SNARE complex. The non-ID region of the syntaxin1 protein was divided into a transmembrane region (residues 265-288), a near-membrane region (residues 258-264) and a SNARE motif (residues 210-257) according to the protein domain, and the SNARE motif was further segmented according to the Munc18-1 binding region (Fig. S8**c**). Regardless of which amino acid sequence of the non-ID region was deleted, SNARE complex droplets did not form (Fig. S8**d**), suggesting that the full-length non-IDR of the syntaxin1 protein is the key region responsible for SNARE phase separation.

**Fig. 5.**
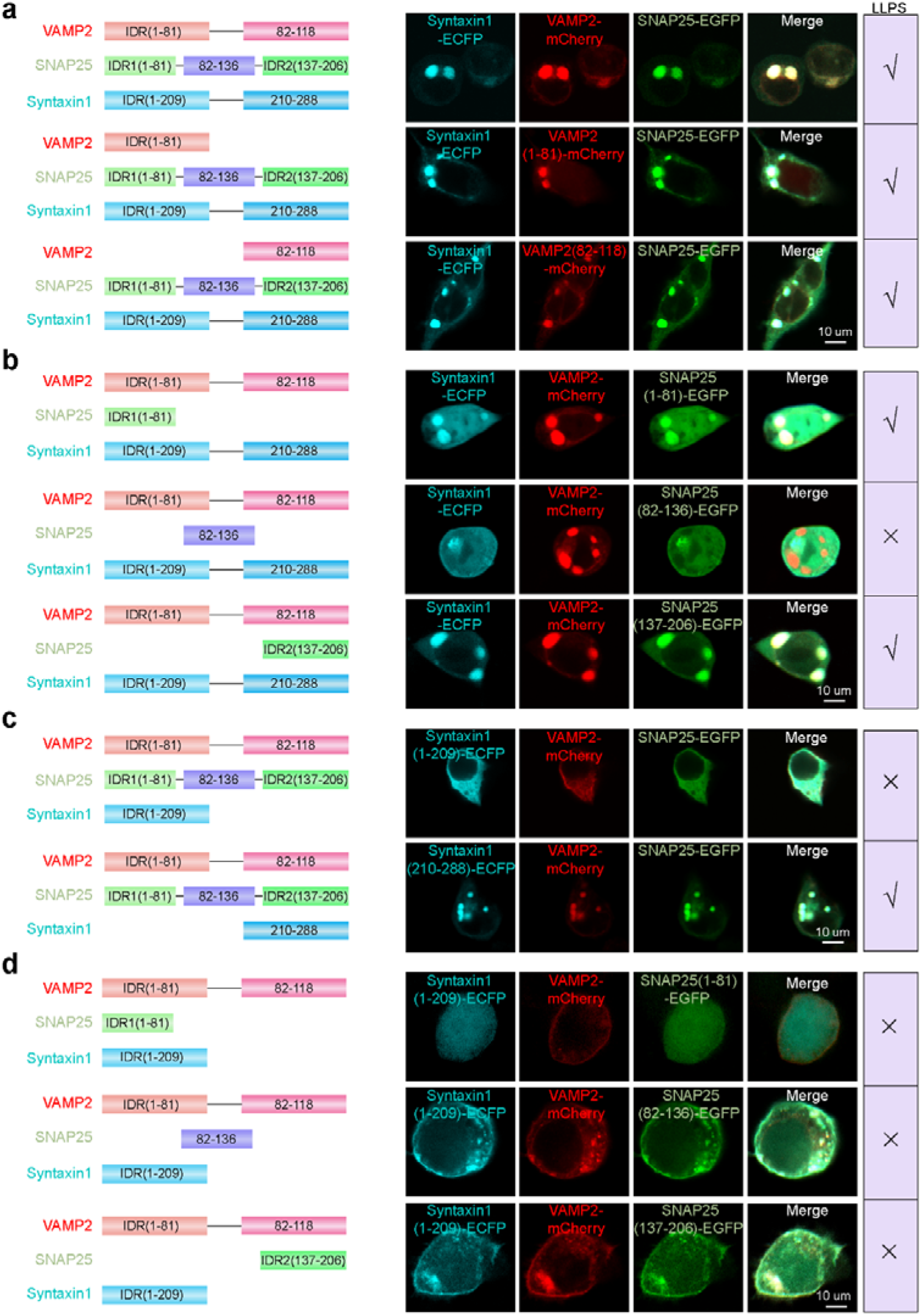
The non-IDRs of syntaxin1 is necessary for the phase separation of the SNARE complex. **a** Full-length IDRs (1-81) or non-IDRs (82-118) of the VAMP2-mCherry construct (1 µg/ml) were cotransfected with syntaxin1-ECFP and SNAP25-EGFP constructs (1 µg/ml) into HEK-293T cells for 24 h, after which images were obtained by confocal microscopy. **b** IDR (1-81), IDR (137-206) or non-IDR (82-136) of the SNAP25-EGFP plasmid (1 µg/ml) was cotransfected with syntaxin1-ECFP and VAMP2-mCherry plasmids (1 µg/ml) into HEK-293T cells for 24 h, after which images were obtained by confocal microscopy. **c** IDR (1-209) or non-IDR (210-288) of the syntaxin1-ECFP plasmid (1 µg/ml) was cotransfected with VAMP2-mCherry and SNAP25-EGFP plasmids (1 µg/ml) into HEK-293T cells for 24 h, after which images were obtained by confocal microscopy. **d** HEK-293T cells were cotransfected with the IDR1 (1-81), IDR2 (137-206) or non-IDR (82-136) of the SNAP25-EGFP plasmid (1 µg/ml) and the IDR (1-209) of the syntaxin1-ECFP and VAMP2-mCherry plasmids (1 µg/ml) for 24 h, after which images were obtained via confocal microscopy. Scale bar: 10 μm.

### SNARE complex phase separation can be regulated by cofactors of fusion machinery

SNARE complex assembly is regulated by numerous presynaptic factors, and SM proteins are of fundamental importance. Munc18-1 and Munc13-1 cooperate to chaperone SNARE assembly. Munc18-1 binds tightly to syntaxin1 folded into a closed conformation, thus impeding the reassembly of the ternary SNARE complex^25^. The MUN domain of Munc13-1 has been found to dramatically catalyze the transition from the closed syntaxin1–Munc18-1 complex to the SNARE complex, possibly by regenerating the SNARE motif of syntaxin1 from the closed conformation and providing a template for SNARE complex formation^26^. To determine the potential effect of Munc18-1 on the phase separation of the SNARE complex, we added increasing amounts of the Munc18-1 plasmid to HEK-293T cells transfected with the mCherry-VAMP2, SNAP25-EGFP, and syntaxin1-ECFP plasmids for 24 h. The size of the SNARE complex droplets gradually decreased with increasing amounts of the Munc18-1 plasmid (Fig. 6**a**, **b**), and this trend was reversed by retransfecting the MUN domain of the Munc13-1 construct, as the droplets size obviously increased with increasing Munc13-1 concentration (Fig. 6 **c** and **d**). These findings indicate that the phase separation of the SNARE complex can be regulated by the major cofactors of the fusion machinery.

**Fig. 6.**
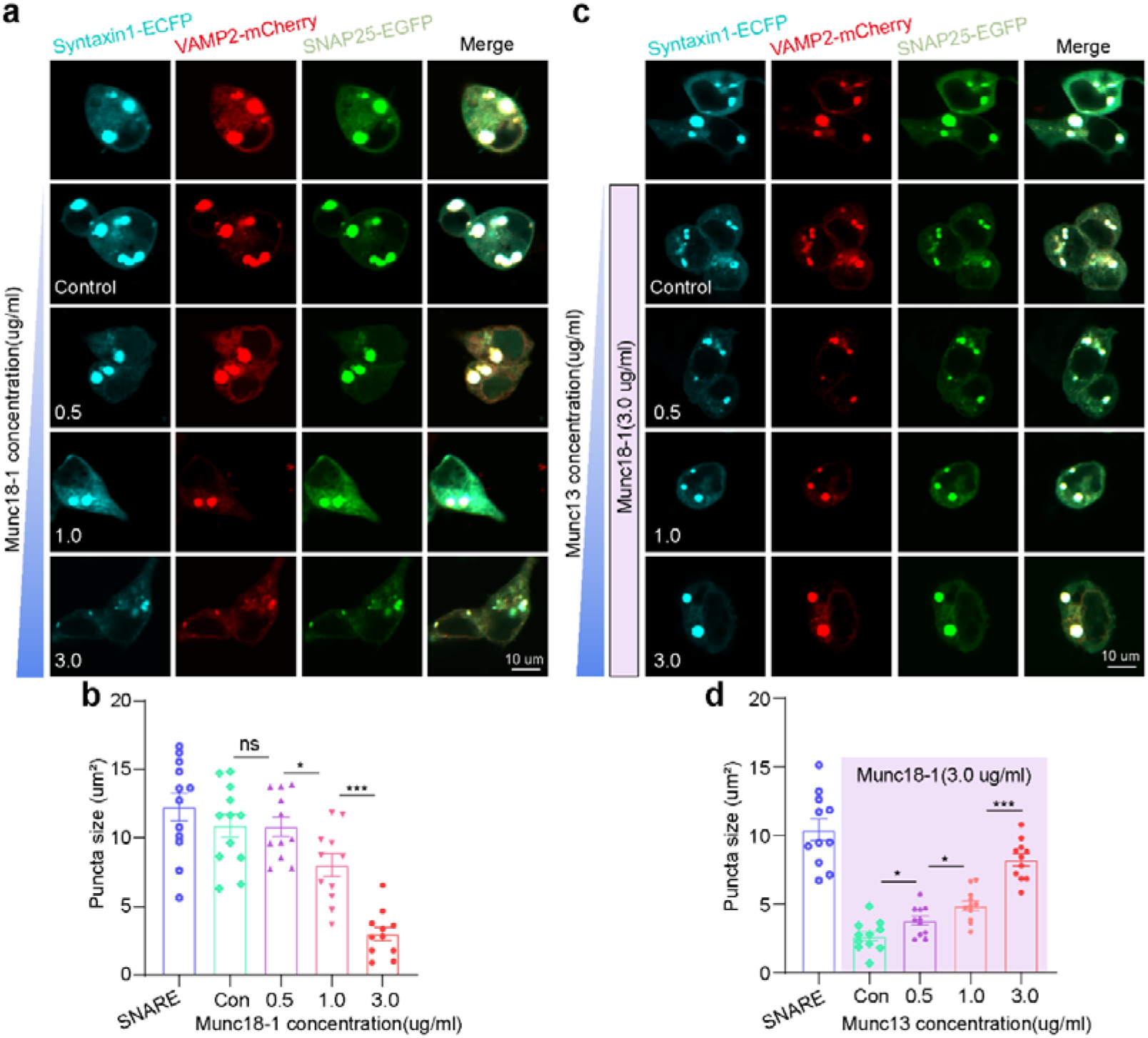
SNARE complex phase separation can be regulated by cofactors of fusion machinery. **a** HEK-293T cells were cotransfected with increasing amounts (0.0, 0.5, 1.0, or 3.0 µg/ml) of the Munc18-1 plasmid and the mCherry-VAMP2, SNAP25-EGFP, or syntaxin1-ECFP plasmid (1.0 µg/ml) for 24 h, after which images were obtained by confocal microscopy under a ×90 objective. **b** Quantification of the puncta size data for the droplets. The average puncta size within cells was quantified, at least 11 cells were randomly selected. The data are displayed as the mean ± SEM. **c** HEK-293T cells were cotransfected with increasing amounts (0.0, 0.5, 1.0, or 3.0 µg/ml) of the Munc domain of the Munc13-1 plasmid and 1.0 µg/ml of the mCherry-VAMP2, SNAP25-EGFP, or syntaxin1-ECFP plasmid and Munc18-1 plasmid for 24 h, after which images were obtained by confocal microscopy under a 90× objective. **d** Quantification of the puncta size data for droplets (n=11). The data are displayed as the mean ± SEM. \**p* < 0.05; \*\**p* < 0.01; \*\*\**p* <0.001.

### SNARE complex phase separation recruits synaptic vesicles in neurons

Next, we investigated whether the SNARE complex can form droplets in neurons. Upon overexpression, droplet formation by the SNARE complex was observed in rat primary neurons (Fig. 7**a**). A GFP construct served as a control and did not form droplets. Since the SNARE complex is the core machinery that governs synaptic transmission, we further explored the role of SNARE complex phase separation in the recruitment of synaptic vesicles. We labeled synaptic vesicles with a synaptophysin antibody and detected obvious colocalization between the SNARE complex phase separation droplets and the synaptic vesicles (Fig. 7**b**), suggesting that SNARE complexes may recruit synaptic vesicles through phase separation and mediate membrane fusion and synaptic transmission.

**Fig. 7.**
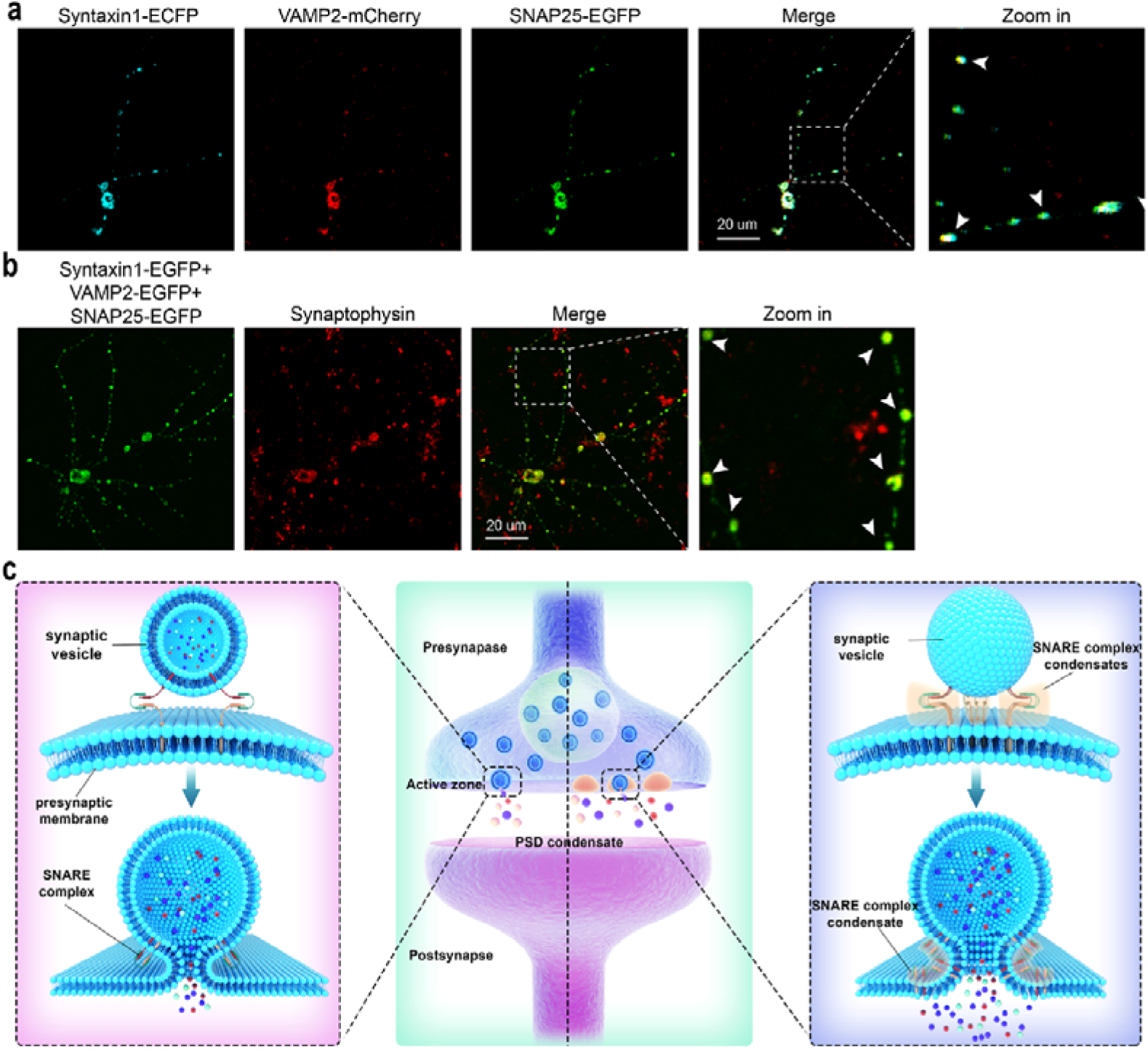
SNARE complex phase separation recruits synaptic vesicles in neurons. **a** Representative fluorescence images of protein droplets in rat primary neurons transfected with syntaxin1-ECFP, VAMP2-mCherry and SNAP25-EGFP plasmids (1 µg/ml) for 24 h. Scale bar, 20 µm. **b** Colocalization of SNARE complex droplets and synaptic vesicles. A Synaptophysin antibody was used to label the synaptic vesicles. Scale bar, 20 µm. **c** Working mode of SNARE complex formation and aggregation driven by phase separation. Left, SNARE complexes mediate membrane fusion processes and neurotransmitter release. Moreover, phase separation mediates the formation and aggregation of the SNARE complex, providing conditions for multiple SNARE complexes to function together, and may be more conducive to overcoming the membrane fusion barrier.

## Discussion

Neuronal communication depends on the exocytosis of neurotransmitters from synaptic vesicles to the synaptic cleft, which is mainly driven by the assembly of the SNARE complexes. However, little is known about the organization pathway of the SNARE complexes and how these SNARE complexes work together to mediate membrane fusion and to ensure sustained neurotransmitter release. In the present study, we found that the formation and aggregation of the SNARE complex are driven by phase separation, which can recruit synaptic vesicles in rat neurons (Fig. 7**c**). Our findings provide new information on the working model underlying the assembly and aggregation of SNARE complexes.

Since the first four-spiral beam structure was established, great progress has been made toward understanding the assembly and disassembly of SNARE complexes. Despite this progress, knowledge on the complex organizational principles of SNAREs remains limited. Neuronal SNARE proteins are considered the engine of the release machinery, as their assembly into the SNARE complex is highly exergonic and produces free energy; this energy pulls the opposing membranes into close proximity and mediates fusion ^27, 2^. According to previous studies, the assembly of one SNARE complex provides a total energy of approximately 85 kBT, including 68 kBT from zippering between VAMP2 and t-SNAREs^28^ and 17 kBT from folding of the t-SNARE complex^29^. This energy is sufficient to overcome the barrier between the synaptic vesicle membrane and the presynaptic membrane and leads to membrane fusion^30, 31^; thus, only one group of the SNARE complex is sufficient to counteract the energy needed for membrane fusion events^32^. However, this perspective is based on the continuous formation of the SNARE complex. In fact, there are multiple cofoctors involved in the regulation of SNARE complex assembly under physiological conditions^33^. Thus, it is almost impossible for a single SNARE complex to be assembled with the conditions necessary for continuous assembly under the precise regulation of these proteins ^34, 35^. This raises a fundamental question: how many SNARE complexes are needed to fuse a vesicle? This critical question has become an areas of focus in the field of membrane fusion research during the past two decades and has yielded inconsistent results.

The number of copies of the SNARE complex needed for membrane fusion is currently under investigation, but the results are inconsistent. Some researchers have estimated that only one SNARE complex can lead to membrane fusion based on single-molecule studies^36^ and liposomes containing a single SNARE molecule with purified synaptic vesicles^32^. Researchers have also shown that three SNARE complexes work simultaneously to support membrane fusion and neurotransmitter release^37^. More studies have suggested that five to eleven groups of SNARE complexes are needed to ensure that fusion pores open and neuronal exocytosis occurs^34, 38, 39, 40, 35, 41, 42^. Much uncertainty remains, mainly due to the lack of effective and direct methods to control and count the number of SNARE complexes related to a fusion event^43^; however, at least several SNARE complexes that work together are needed to ensure membrane fusion and sustained neurotransmitter release in vivo. Another vital question arises: does the SNARE complex function as multiple copies that are dispersed or clustered together? If the latter is true, what forces hold the SNARE complexes together?

Extensive evidence in recent years has revealed that LLPS is likely a ubiquitous organization principle that drives the formation of nonmembrane-bound organelles^44^. Notably, in recent years, it has been established that phase separation of active zone scaffold components may contribute to synaptic vesicle tethering, docking, priming, and even short-distance vesicle transport process^22^. How about the fusion process? In vitro reconstitution studies have demonstrated that membrane fusion can be catalyzed by the SNARE complex without any other partners, but the duration of this process is too long^45^. Recent reconstitution fusion assays in the presence of SNARE complex regulatory factors, including Munc18-1, Munc13-1, NSF, a-SNAP, complexin, and synaptotagmin, have clearly increased the fusion speed to the millisecond level^8, 46^. Can phase separation further accelerate the membrane fusion process? Several favorable observations support the possibility that the SNARE complex undergoes phase separation. First, SNARE proteins are among the most abundant proteins in presynaptic boutons, with more than 100 µM in each bouton^47^. This concentration is much greater than the number of SNARE proteins needed for membrane fusion events^48^. Zhang’s group hypothesized that if the SNARE complex and regulators are enriched by phase separation in a manner similar to synaptic vesicle clustering mediated by synapsin, membrane fusion events can further accelerate to physiologically relevant rates^49^. In fact, several important regulatory proteins of the SNARE complex, such as α-synuclein and synaptotagmin1, have recently been reported to undergo phase separation and regulate synaptic secretion through phase separation^19, 50^. Second, SNARE proteins are not uniformly distributed in the membrane. Instead, syntaxin1 and SNAP25 are concentrated in clusters within the presynaptic membrane that define docking and fusion sites for exocytosis^51, 52^, and each synaptic vesicle harbors approximately 70 copies of active VAMP2 molecules^53^, ensuring that multiple SNARE proteins are always available. Third, spontaneous aggregation of SNARE protein monomers can be observed at the recombinant protein level in vitro, and SNARE proteins contain extensive IDR structures, implying that phase separation may occur in the SNARE complex. In the present study, we determined that the SNARE complex can undergo phase separation by performing a FRAP assay, droplet fusion or division assay, and inducing interference with phase separation-specific inhibitors; this process relies on the non-IDRs of the syntaxin1 protein. Notably, SNARE complex phase separation has the ability to recruit synaptic vesicles in rat neurons, suggesting that phase separation of SNARE complexes may function in synaptic transmission. It has been demonstrated that the number of copies of SNARE complexes per vesicle affects the speed of Ca^2+^-dependent transmitter release^33^. Since phase separation ensures that multiple SNARE complexes work together, both the number of SNARE complexes and the energy needed to overcome the energy barrier for efficient synaptic membrane fusion were confirmed. Hence, the phase separation-driven SNARE complex cluster model can effectively trigger membrane fusion and neurotransmitter release. Thus, it was determined whether the SNARE complex functions as a single copy or as a cluster and the question of how many SNARE complexes are needed to work together to maintain sustained neurotransmitter release is properly answered.

When defining the phase separation of the SNARE complex, we detected partial but incomplete colocalization between the droplets and the endoplasmic reticulum during the exclusion of membranous organelles. This is reasonable because the endoplasmic reticulum is an important organelle for protein synthesis^54^. This finding indirectly indicates that these droplets are protein condensates. Notably, IDRs, especially those containing a low-complexity domain, have been reported to drive phase separation^13^. However, we found that the phase separation of the SNARE complex does not depend on the IDR of the syntaxin1 protein, possibly because the IDRs of these proteins mainly confer an additional role; the proteins primarily span a radius greater than that of a folded region, thus increasing the likelihood that binding partners are recruited^55, 56^.

Following vesicle fusion, SNARE complexes is disassembled by N-ethylmaledimide-sensitive factor (NSF) via ATP hydrolysis with the help of soluble NSF attachment protein (SNAP), which extracts SNARE proteins into individuals for another round of membrane fusion^57^. The assembly disassembly cycles of SNARE complexes are also regulated by many key factors. Here, by employing FRAP and droplet fusion or division assays, we observed the dynamic properties that underly the phase separation of SNARE complex. Through the dynamic nature of phase separation, the SNARE complex is sensitive to the peripheral environment; thus, the complex can be regulated by numerous cofactors and gains its recycling properties. Our data suggest that phase separation of the SNARE complex is sensitive to general regulators of phase separation, such as the salt environment, crowding agents, and protein concentration, and is also regulated by cofactors of fusion machinery, such as Munc18-1 and Munc13-1. Notably, the molecular weight of the fluorescent proteins employed in this study is equivalent to or greater than that of SNARE proteins, which may seriously affect the dynamics of SNARE complex phase separation and exchange with surrounding materials; therefore, it is reasonable that the actual dynamics of SNARE complex phase separation are much greater than those observed experimentally. In subsequent experiments, labeling SNARE proteins with more suitable fluorescent tags or probes may better reflect the dynamic characteristics of the SNARE complex.

There are several limitations in this study. First, although we used strict controls, the effect of tag proteins on the phase separation of the SNARE complex cannot be completely ruled out. Second, whether the SNARE complex undergoes phase separation in vivo and how phase separation contributes to membrane fusion and synaptic transmission remain unclear. Moreover, the interactions between SNARE complex phase separation, other membrane fusion cofactors and presynaptic phase condensates, especially those located in the presynaptic active zone, are unknown. In addition, since membrane fusion is a universal process in eukaryotic cells, it would be interesting to determine whether the SNARE fusion machinery mediated by phase separation also affects other cellular membrane fusion processes that are determined by SNARE proteins, such as mitochondria fusion^58^, autophagosome-lysosome fusion^59^, endosomal and phagosomal trafficking^60^, and yeast vacuole fusion^61^.

## Conclusions

Our findings demonstrates that phase separation is a promising way to mediate the formation and aggregation of the SNARE complex, and further identified that the non-IDRs of syntaxin1 is necessary for the phase separation of the SNARE complex. Our work answers an vital scientific question: does the SNARE complex function as multiple copies that are dispersed or clustered together to ensure sustained neurotransmitter release. In sum, phase separation provides an ideal working model for SNARE complex-mediated membrane fusion and neurotransmitter release.

## Abbreviations

SNARE: soluble *N*-ethylmaleimide-sensitive factor attachment protein receptor
SM: Sec1/Munc18
LLPS: liquidLliquid phase separation
FRAP: fluorescence recovery after photobleaching
IDRs: Intrinsically disordered regions 1,6-HD 1,6-Hexanediol
NSF: N-ethylmaledimide-sensitive factor
SNAP: soluble NSF attachment protein

## Acknowledgements

Not applicable.

## Author contributions

Y.W., L.C., and L.D. designed the experiments; F.C., Q.K., S.F., H.C., Q,H., F.M, Y.C., S.L., W.W., P.X., Y.S., J.C., L.T., and X.S. performed research; F.C., Q.K., and S.F. analyzed data; F.C. wrote the paper; L.C. and Y.W. revised the paper. All authors have read and approved the final version of manuscript.

## Availability of data and materials

All data generated and analyzed during this study are either included in this article or are available from the corresponding author on reasonable request.

## Declarations

### Ethics approval and consent to participate

All animal experimental procedures were approved by the Laboratory Animal Ethics Committee of Guangdong Medical University.

### Consent for publication

Not applicable.

## Supplementary Information

**Fig. S1.**
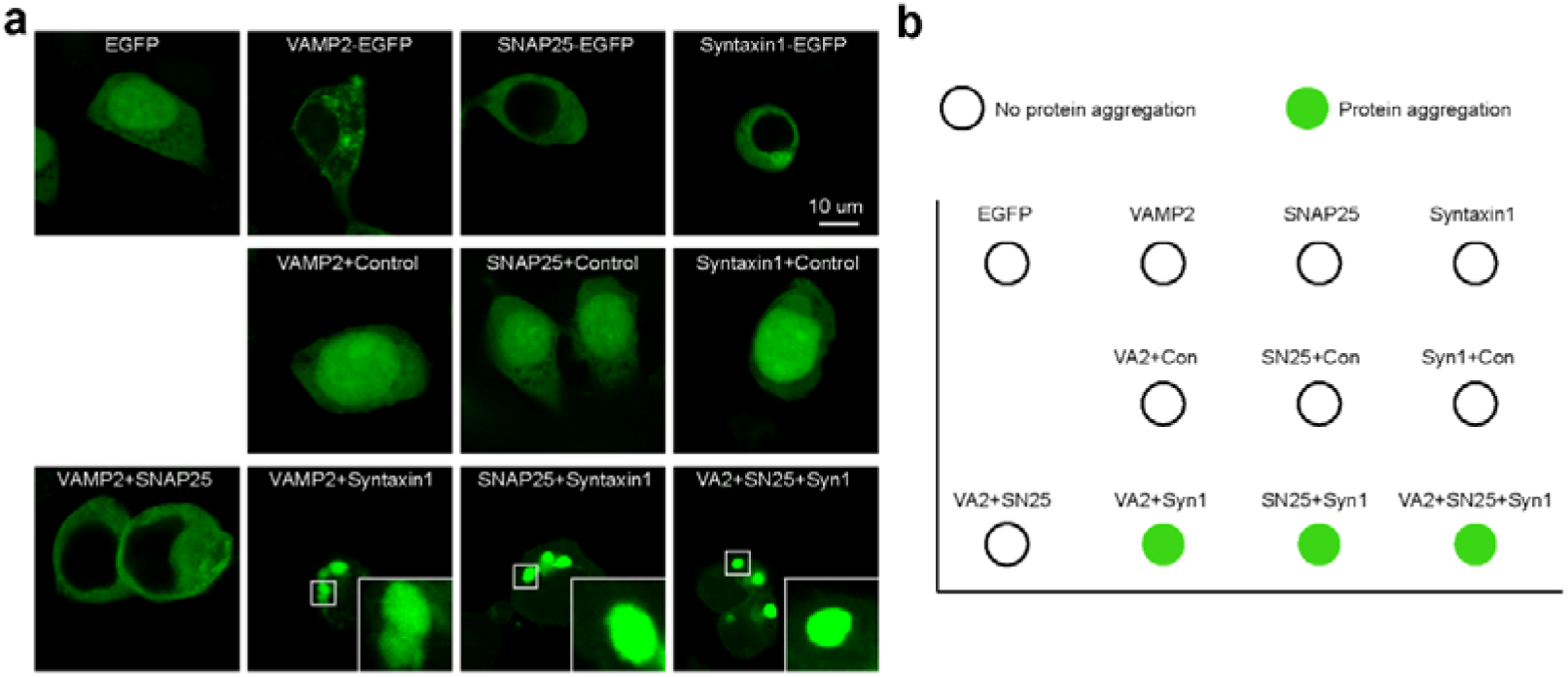
The SNARE complex forms protein condensates in HEK-293T cells. **a** Representative images of HEK-293T cells transfected with 1 µg/ml EGFP, VAMP2-EGFP, SNAP25-EGFP or syntaxin1-EGFP plasmid alone; transfected with 1 µg/ml EGFP and VAMP2-EGFP, EGFP and SNAP25-EGFP, or EGFP and syntaxin1-EGFP plasmids; and transfected with VAMP2-EGFP and SNAP25-EGFP, VAMP2-EGFP and syntaxin1-EGFP, SNAP25-EGFP and syntaxin1-EGFP, or VAMP2-EGFP and SNAP25-EGFP and syntaxin1-EGFP plasmids for 24 h. (scale bar, 10 µm). **b** Regime diagram illustrating the protein condensates shown in Fig. S1a.

**Fig. S2.**
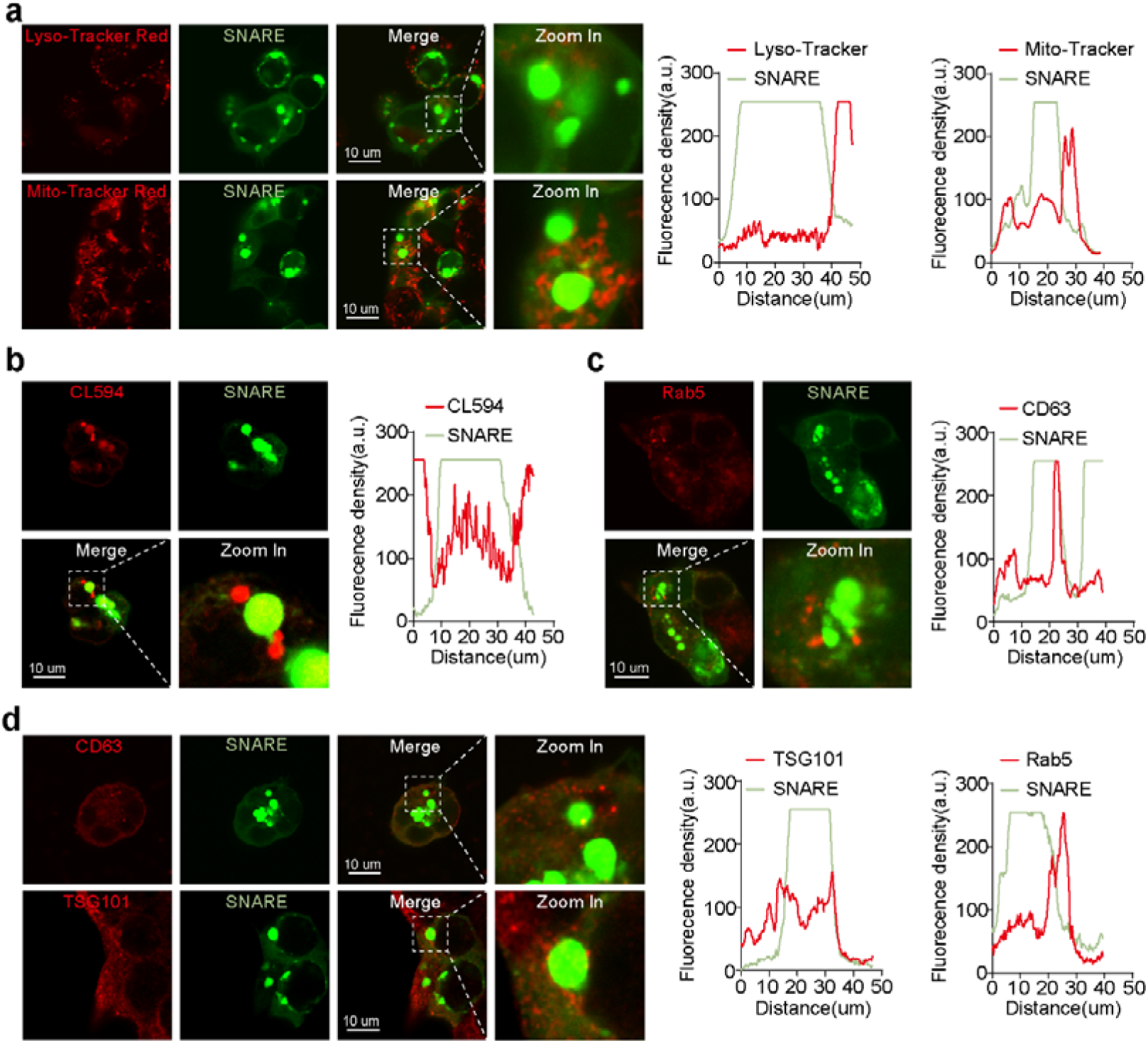
SNARE complex droplets do not colocalize with membrane organelles, exosomes or vesicles. **a** HEK-293T cells were transfected with 1 µg/ml VAMP2-mCherry, SNAP25-EGFP and syntaxin1-ECFP plasmids for 24 h, and representative images of droplets colocalization with lysosomes (LysoTracker Red) and mitochondria (Mito-Tracker Red) were obtained by confocal microscopy. (scale bar, 10 µm). **b** Representative images showing the colocalization of SNARE droplets with the endoplasmic reticulum marker CL594. (scale bar, 10 µm). **c** Representative images showing the colocalization of SNARE droplets with the vesicle marker Rab5. (scale bar, 10 µm). **d** Representative images of SNARE droplets colocalization with the exosomal markers CD63 and TSG101. (scale bar, 10 µm).

**Fig. S3.**
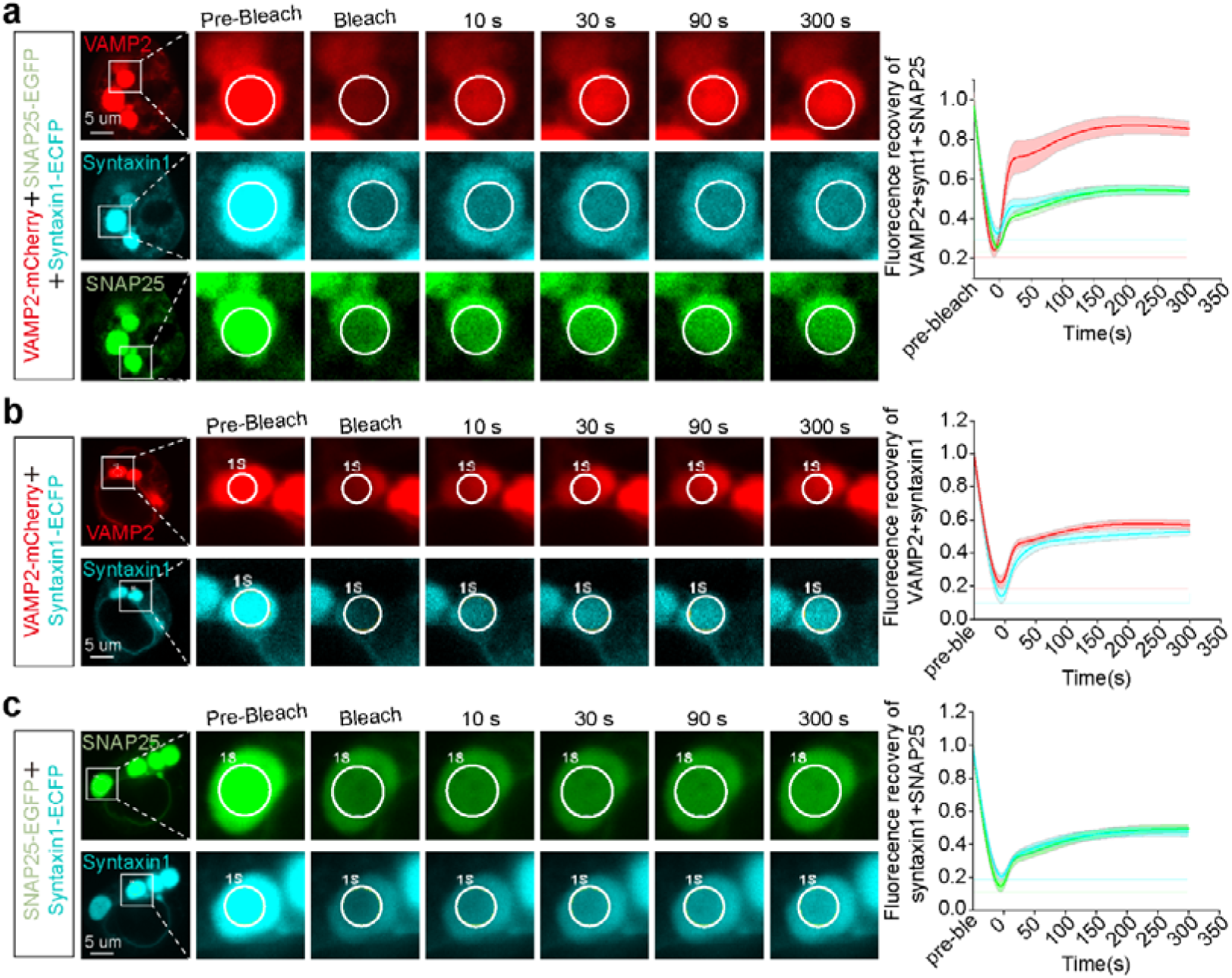
FRAP assay was performed to examine the kinetics of the SNARE complex in cells. Representative images from live-cell imaging of VAMP2-mCherry, SNAP25-EGFP and syntaxin1-ECFP constructs (**a**), VAMP2-mCherry and syntaxin1-ECFP constructs (**b**), and SNAP25-EGFP and syntaxin1-ECFP constructs (**c**) overexpressed in HEK-293T cells (1 µg/ml for 24 h) before and after photobleaching. White circles indicate bleached regions. Scale bar: 5 μm. Six droplets were used for FRAP analysis.

**Fig. S4.**
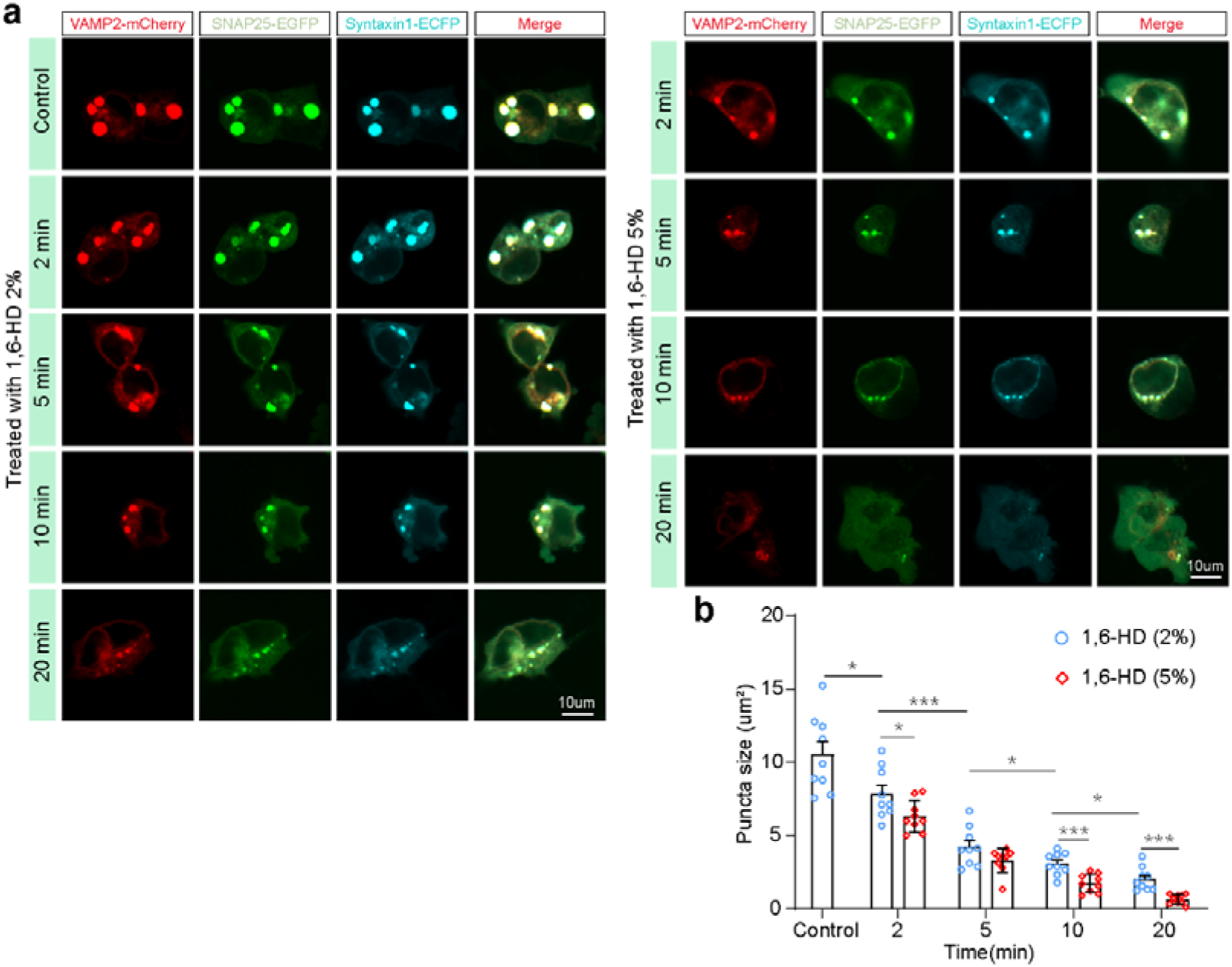
Effects of 1,6-HD on the condensates formation of the SNARE complex. **a** HEK-293T cells were cotransfected with VAMP2-mCherry, SNAP25-EGFP and syntaxin1-ECFP constructs (1 µg/ml) for 24 h and then exposed to different doses (0, 2%, or 5% volume ratio) of 1,6-HD for various durations (0, 2, 5, 10, or 20 min), after which images were obtained via confocal microscopy. Scale bar: 10 μm. **b** Quantification of the average puncta size data for droplets within cells (n=9). The data are displayed as the mean ± SEM. \**p* < 0.05; \*\**p* < 0.01; \*\*\**p* <0.001.

**Fig. S5.**
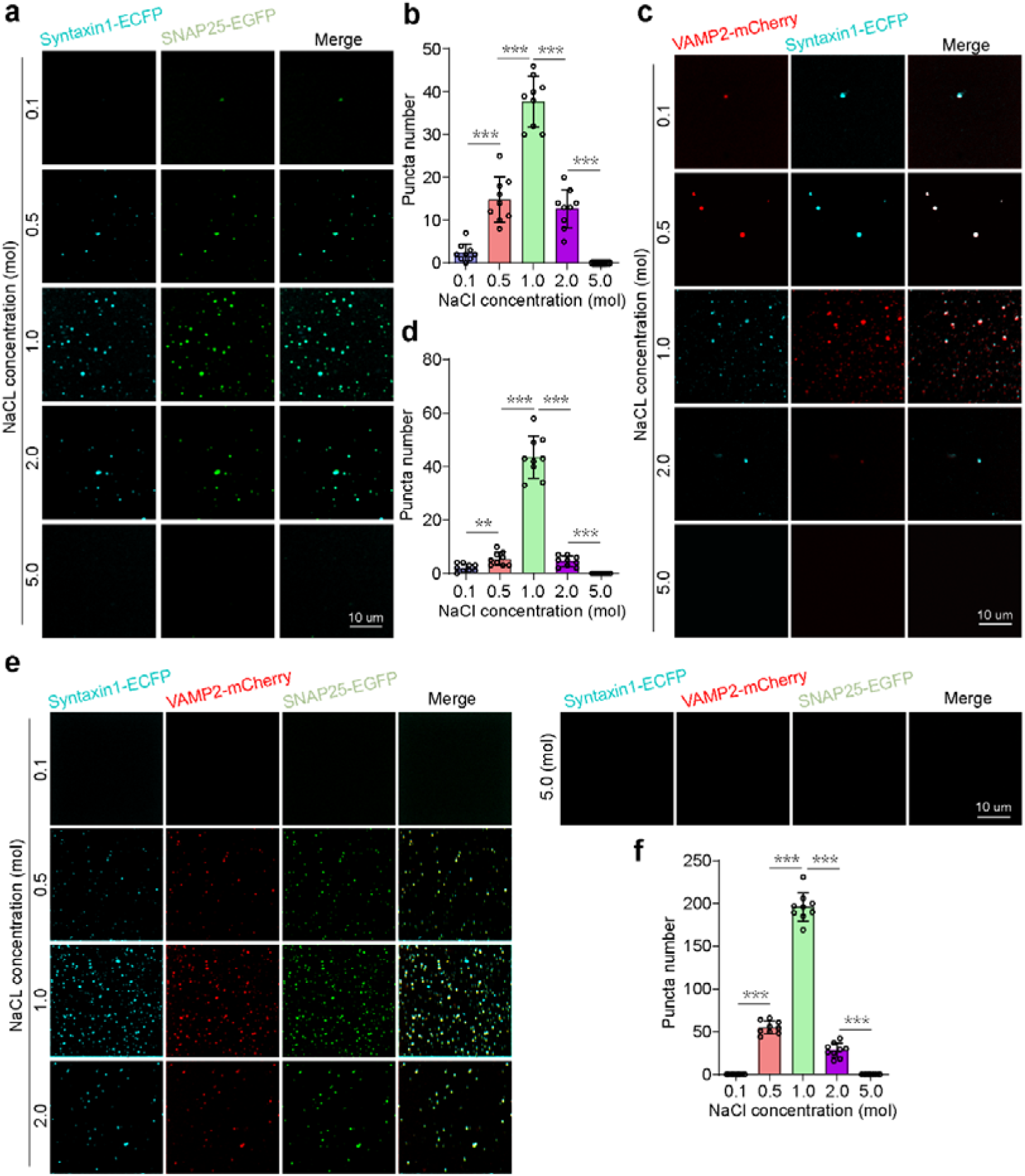
SNARE complex phase separation is sensitive to saline ions. Syntaxin1-ECFP and SNAP25-EGFP fusion proteins (**a**, **b**), Syntaxin1-ECFP and VAMP2-mCherry fusion proteins (**c**, **d**), and Syntaxin1-ECFP, SNAP25-EGFP and VAMP2-mCherry fusion proteins (**e**, **f**) were subjected to different doses of NaCl (0.1, 0.5, 1.0, and 2.0 mol) in buffer containing crowding reagent (PEG-8000, 10% w/v) for 10 min, and images were obtained by confocal microscopy. Nine fields of view were randomly selected for statistical analysis. The data are presented as the mean ± SEM. Scale bar: 10 μm. \**p* < 0.05; \*\**p* < 0.01; \*\*\**p* <0.001.

**Fig. S6.**
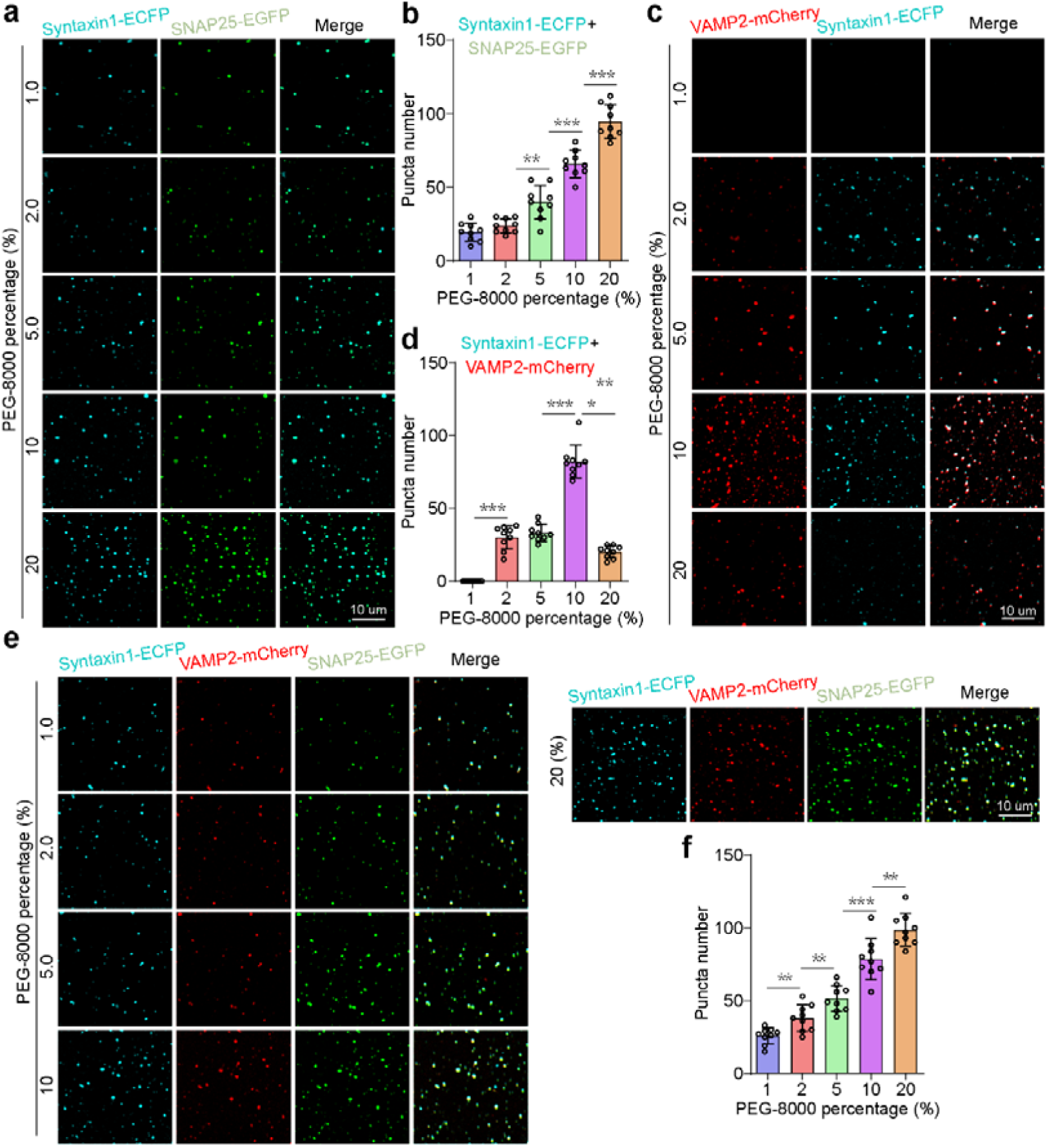
SNARE complex phase separation was enhanced by the addition of a crowding reagent. Syntaxin1-ECFP and SNAP25-EGFP fusion proteins (**a**, **b**), Syntaxin1-ECFP and VAMP2-mCherry fusion proteins (**c**, **d**), and Syntaxin1-ECFP, SNAP25-EGFP and VAMP2-mCherry fusion proteins (**e**, **f**) were subjected to different doses of NaCl (0.1, 0.5, 1.0, and 2.0 mol) in buffer containing crowding reagent (PEG-8000, 10% w/v) for 10 min, and images were obtained by confocal microscopy. Nine fields of view were randomly selected for statistical analysis. The data are presented as the mean ± SEM. Scale bar: 10 μm. \**p* < 0.05; \*\**p* < 0.01; \*\*\**p* <0.001.

**Fig. S7.**
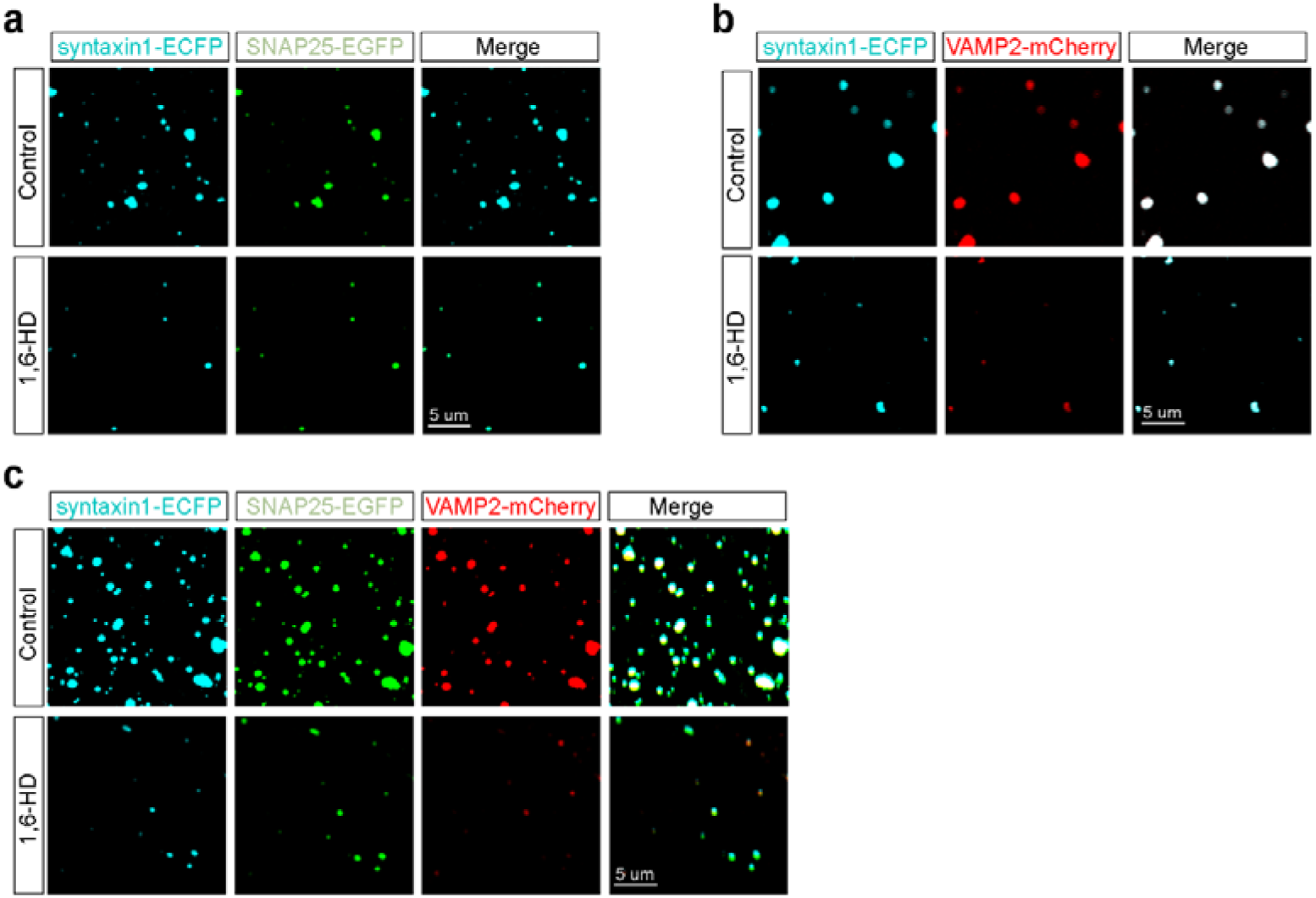
Effects of 1,6-HD on the condensates formation in vitro studies with purified proteins. Representative images of syntaxin1-ECFP and SNAP25-EGFP (G), syntaxin1-ECFP and VAMP2-mCherry (H), and syntaxin1-ECFP, SNAP25-EGFP and VAMP2-mCherry droplets before and after treatment with 1,6-HD. Scale bar, 5 μm.

**Fig. S8.**
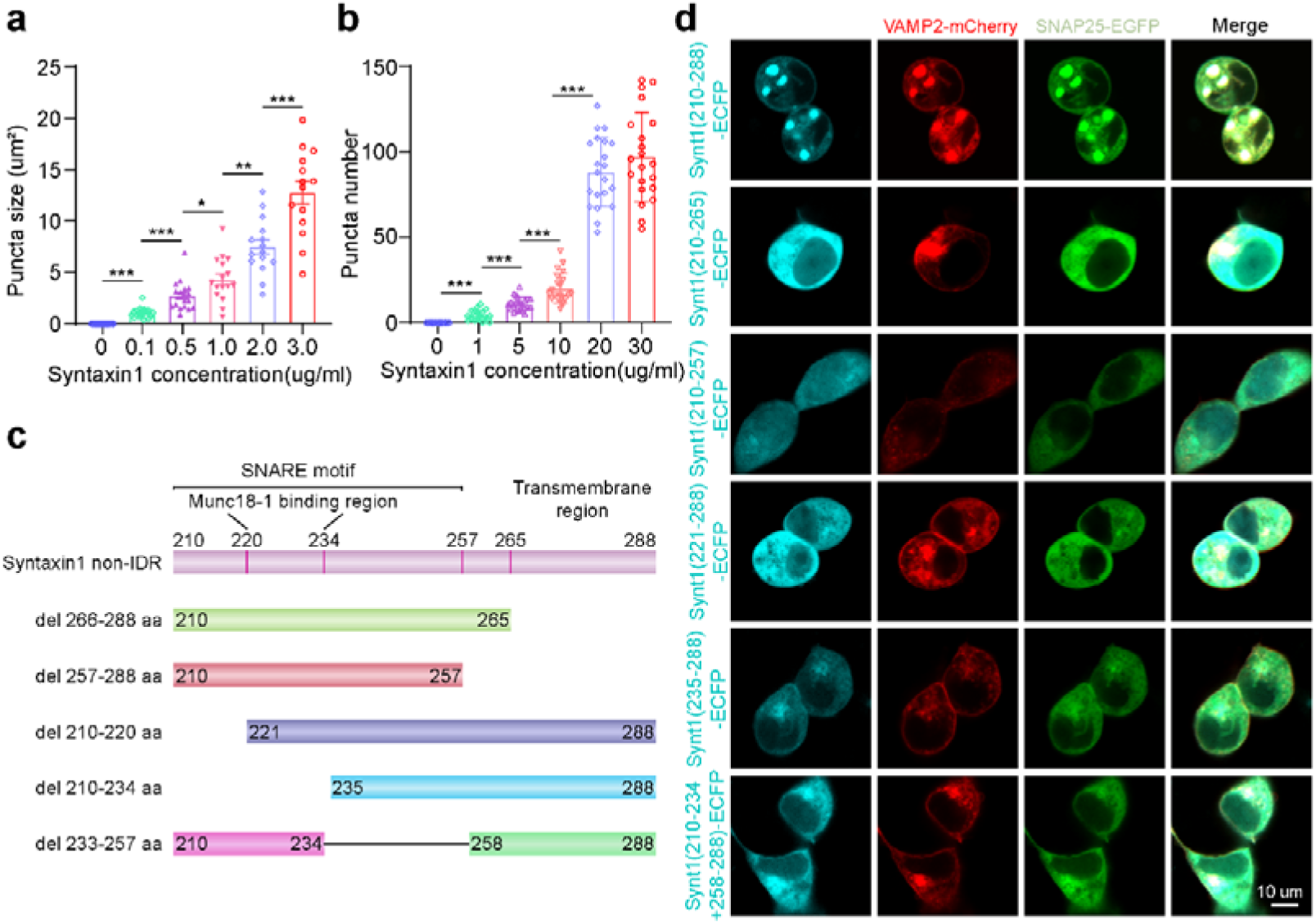
The IDR of syntaxin1 is not responsible for the phase separation of the SNARE complex. **a** Quantification of the average puncta size within cells of Fig. 4a. Six cells were chosen for the control group, and at least 14 cells were included in the other groups. The data are displayed as the mean ± SEM. **b** Quantification of the droplets number (n≥20) of Fig. 4b. The data are displayed as the mean ± SEM. \**p* < 0.05; \*\**p* < 0.01; \*\*\**p* <0.001. **c** Domain schematic and constructs used in the experiment. The non-IDRs (residues 210-288) of the syntaxin1 protein were divided into transmembrane regions (residues 265-288), near-membrane regions (residues 210-288) and SNARE motifs (residues 210-288) according to the protein domain. The SNARE motif was further divided into three parts (residues 210-219, 220-234, and 235-257) according to the binding region of the Munc18-1 protein. **d** The above five regions of the non-ID region of the syntaxin1 protein were deleted, and the remaining regions were cotransfected with the VAMP2-mCherry and SNAP25-EGFP plasmids. Images were obtained by confocal microscopy after 24 h of transfection. Scale bar: 10 μm.

